# Cocaine Blocks Cholinergic Activity in the Medial Habenula Prior to But Not After Induced Preference for the Drug

**DOI:** 10.1101/2025.03.20.644247

**Authors:** Yousheng Jia, Chenyi C. Yang, Julie C. Lauterborn, Christine M. Gall, Marcelo A. Wood, Gary Lynch

**Affiliations:** Dept. of Anatomy and Neurobiology, University of California, Irvine, 92697, USA; Dept. of Neurobiology and Behavior, University of California, Irvine, 92697, USA.; Dept. of Psychiatry and Human Behavior, University of California, Irvine, 92868, USA

**Keywords:** acetylcholine, dopamine, drug abuse, addiction, nicotinic

## Abstract

Descending projections from the medial habenula potently influence brainstem systems associated with reward and mood. Relatedly, the ventral, cholinergic segment of the nucleus has been linked to nicotine and cocaine addiction. Here we report that cocaine has no effect on baseline firing in the ventral medial habenula but entirely blocks the self-sustained activity initiated by endogenous acetylcholine. This effect was not altered by antagonists to dopamine receptors and thus presumably reflects a direct action on cholinergic receptors. Remarkably, cocaine had no effect on endogenous cholinergic activity in mice that had been extinguished from an induced cocaine preference. In all, the drug has potent effects, albeit through an exotic mode of action, on the medial habenula and these are eliminated by prior experience with the drug. These results describe a novel target for cocaine that is plausibly related to the psychological effects of the drug, and an unexpected consequence of earlier use.

## Introduction

Cocaine is a stimulant drug that among other effects induces a sense of euphoria. Its major neurobiological action is to block dopamine transporters and thereby elevate intra-synaptic dopamine levels.^1-4^ Among other consequences, these effects enhance transmission in the projections from the ventral tegmental area (VTA) of the midbrain to the ventral striatum (nucleus accumbens), a connection that constitutes a primary reward system of the brain.^5-7^ In addition to its established dopamine targets, there is evidence indicating that cocaine acts on nicotinic acetylcholine receptors (nAChRs) including those containing α4β2^4, 6^ and α3β4^8-10^ subunits. Notably, α4β2 nAChRs are highly expressed in the cell bodies and terminals of dopaminergic neurons.^11-13^ Other studies indicate that nAChRs affect release kinetics for dopamine terminals.^14-17^ These results suggest that cocaine’s effects on dopamine-mediated reward are modulated by the drug’s actions on cholinergic receptors. In support of this, recent studies have provided direct evidence that cocaine influences activity-driven dopamine release via nAChRs in the ventral striatum.^18^

Little attention has been given to the possibility that cocaine affects motivation and reward via nicotinic binding outside of the dopaminergic system. Dopamine neurons in the VTA are innervated by cholinergic neurons in the midbrain^19, 20^ and indirectly, via the interpeduncular nucleus, in the medial habenula (MHb). ^21, 22^ The latter epithalamic region is of particular interest because it includes a ventral division (vMHb) that contains exceedingly dense concentrations of cholinergic neurons^23, 24^ and nicotinic receptors, including marked expression of the α3 and β4 subunits,^25 26^ and with the interpeduncular nucleus has been linked to mood control.^27-29^ Notably, cocaine has been reported to bind and inhibit α3β4 nAChRs^10, 30^ and high affinity ligands for these receptors have been shown to attenuate cocaine-induced conditioned place preference.^31^ Taken together, these observations suggest a hypothesis in which cocaine, while exerting its primary effect on dopamine release and reuptake,^1, 32^ also inhibits activity in the MHb and thereby relaxes a constraint on dopamine neurons. The present studies tested the prediction that cocaine has a potent effect on ACh-driven neuronal spiking in the MHb.

Repeated use of cocaine results in a strong craving for the drug, an addiction-like effect that gradually decreases over an extended recovery period.^33-35^ Unfortunately, subsequent contact commonly causes a rapid reinstatement (‘relapse”) of the addicted state.^36, 37^ Much has been learned about the cellular and genomic changes associated with cocaine dependency^3,38-40^ but little is known about the neurobiological factors responsible for relapse. However, and relevant to the above comments, preclinical work has shown that renewed usage activates the vMHb and that this is critical for relapse.^37^ Related studies confirmed that repeated administration of cocaine results in epigenetic changes in the nucleus that play a central role in subsequent reactivation of drug seeking by mice. ^41, 42^ These findings raise the possibility that repeated cocaine usage produces lasting changes in the cholinergic system of the vMHb and that this alters the normal response of the nucleus to acute treatment with the drug. If so, then the effects of cocaine on ACh-mediated neuronal activity as observed in naïve mice should be markedly different in animals that had recovered from drug craving. Tests of this argument were conducted in the work reported here.

## Methods

### Animals

Adult 2.5-3 month old male C57BL/6J mice from an in-house colony or ChAT-IRES-Cre mice (Jackson Laboratory; stock # 006410) were bred and used in the present work. All mice were maintained on a 12-hour light/dark cycle with food and water provided *ad libitum*. Work was conducted in accordance with the National Institutes of Health guideline for animal care and use, and experiments were approved by the Institutional Animal Care and Use Committee of the University of California, Irvine.

### Treatments and behavior

For some experiments, animals received cocaine in a version of the conditioned place preference (CPP) paradigm.^37, 43, 44^ The testing apparatus was a rectangular acrylic box (60 cm × 40 cm × 20 cm) with three chambers, and doorways between the middle and side chambers. Each side chamber had distinct visual cues on the walls. All behavior was recorded with an overhead camera for subsequent scoring off-line. On days 1-3 of the cocaine CPP paradigm, mice (n = 16) were handled for 2 min each day and returned to their home cages. On day 4, each mouse was placed in the 3-chamber apparatus and allowed to freely explore for 15 min. Time spent (in seconds) in each side chamber was measured. On days 5-8, mice were given cocaine-HCl (10 mg/kg, IP; Sigma Aldrich, # C5776) and 0.9% saline on alternating days. On days 5 and 7, mice that received cocaine were placed in same side chamber immediately following the injection whereas on saline injection days 6 and 8 they were placed in the opposite side chamber; the pairing of cocaine/Sal to the chambers was counter balanced across animals. In all cases, mice were restricted to the side chamber for 30 min before being returned to their home cages. On day 9, the mice were allowed 15 min to re-explore the 3-chamber apparatus and time spent in each side chamber was measured. Greater time spent in the cocaine-paired chamber was used as an indication of place preference. On days 10-14 for the ‘extinction’ phase, the animals were allowed to freely explore the apparatus for 15 min each day. Finally, for evaluation of reinstatement of place preference, the mice were randomly divided into two equal sized groups (n = 8 each), one of which received a saline injection before being placed in the test apparatus and the other a cocaine injection (5 mg/kg, IP). The animals were then placed in the central compartment and allowed to freely explore all three chambers for 15 min. In summary, all mice were treated the same until day 15 and then were subdivided into saline vs. cocaine groups. A preference score (seconds) was calculated for each phase of the paradigm: time spent in the cocaine-paired chamber minus time spent in the saline-paired chamber. For some analyses of behavior prior to reinstatement (day 15), paired Student’s t tests were used. For other analyses, within-group comparisons were made using two-way RM-ANOVA followed by Tukey’s multiple comparison test for all 15 days for each group (that is, an animal’s condition on day 15 was used to define its group for days 3-14). All statistics were conducted using Prism (version 10.3.1) and mean ± standard error values are reported.

For electrophysiological studies, mice went through the first 14 days of the paradigm and were used for recordings one day later; these animals did not receive any additional drug following the conditioning period.

### AAV-injections

For transfection of MHb cholinergic neurons, 0.5 µL of either AAV1-hSyn-DIO-GFP (7 x 10^12^ vg/ml; Addgene, # 50457) or AAV2.8-hSyn-DIO-mCherry (5.3 × 10^12^ vg/ml; UNC Vector Core) was injected into the MHb bilaterally (from bregma: M/L, ±0.35 mm; A/P, −1.5 mm; D/V; −3.0) as previously described.^37^ For transfection of septal cholinergic neurons, ChAT-Cre mice received two 0.5-μL injections of AAV1-EF1a-DIO-hChR2(H134R)-eYFP-WPRE-hGH (7 x 10^12^ vg/ml; Penn Core) into medial septum (from bregma: M/L, midline for both; A/P +1.3 and D/V -4; A/P +0.7 and D/V -3.9). In each case animals were allowed to recover a minimum of 4 weeks prior to further experimental use.

### Electrophysiology

Coronal slices through the diencephalon were prepared as described.^37^ Briefly, mice were anesthetized by isoflurane inhalation and decapitated. The brain was trimmed, and coronal slices (270 µm thick) containing the MHb were cut using a vibrating microtome (Leica VT1000s). Slices were cut in a choline-based solution containing the following (in mM): 110 choline chloride, 2.5 KCl, 1.25 NaH_2_PO_4_, 0.5 CaCl_2_, 7 MgSO_4_, 26 NaHCO_3_, 25 glucose, 11.6 sodium ascorbate, and 3.1 sodium pyruvate at room temperature. After slice cutting, sodium chloride was progressively spiked into the choline solution every 5 min for 20 min at 32°C, as described.^45^ The slices were allowed to recover for at least an additional 30 min at room temperature in artificial cerebrospinal fluid (ACSF) prior to recording; ACSF contained (in mM): 124 NaCl, 3 KCl, 1.25 KH_2_PO_4_,1.5 MgSO_4_, 26 NaHCO_3_, 2.5 CaCl_2_, and 10 glucose (300–310 mOsm, pH 7.4).

All solutions were saturated with 95% O_2_ and 5% CO_2_. The slices were then placed in a recording chamber, submerged, and continuously perfused at 2–3 ml/min with oxygenated ACSF. Loose cell attached (40-50 MΩ) and whole-cell recordings were achieved from single neurons in the vMHb using patch-clamp amplifiers (Axopatch 200A) under infrared differential interference contrast. Electrical stimulation was delivered by a bipolar tungsten stimulation electrode (World Precision Instruments) placed >50 µm from the recorded neuron. All recordings were performed at 32 ± 1 °C using an automatic temperature controller (Warner Instruments). The loose cell attached recording glass pipettes (5-7MΩ) were filled with 0.9% saline. Whole-cell recordings were made with 4–7 MΩ recording pipettes filled with a solution containing (in mM): 80 KMeSO_3_, 60 KCl, 10 HEPES, 0.2 EGTA, 2 QX-314, 2 Mg-ATP, 0.3 Na-GTP with osmolarity adjusted to 290– 295 mOsm and pH 7.4. Excitatory postsynaptic currents (EPSCs) were recorded by clamping the cell at −70 mV in the presence of 50 μM APV (Hello Bio, HB0225) and 20 μM DNQX (Hello Bio, HB0261). Data acquisition and analysis were performed using DigiData 1550A digitizers and analysis software pClamp 10 (Molecular Devices). Signals were filtered at 2 kHz and sampled at 5 kHz. Drugs were infused for 20 minutes after a 10-minute baseline period. Physostigmine alone was delivered to two groups of slices (n= 6 each) at different stages of the study and the resulting data used for comparison with each of the other drug conditions. For statistical analyses of neuronal firing, mean firing rates were compared for baseline vs. the last five minutes of drug infusion.

Drugs and concentrations used: cholinesterase inhibitor physostigmine hemisulfate (Tocris Bioscience, # 0622), 10 µM; selective D_2_-like antagonist sulpiride (Tocris Bioscience, # 0895) 2.5 µM; selective D_1_-like antagonist SCH23390 hydrochloride (Tocris Bioscience, # 0925), 2.5 µM; non-competitive nicotinic ACh receptor antagonist mecamylamine hydrochloride (Tocris Bioscience, # 2843), 10 µM; cocaine hydrochloride (Sigma Aldrich, # C5776), 10 µM; GABAA receptor antagonist bicuculline (Tocris, # 0130), 20 µM.

### Immunolabeling of AAV-expressing cholinergic neurons

For visualization of transfected MHb cholinergic neurons, coronal slices through the habenula of AAV-expressing ChAT-Cre mice were prepared on a vibrotome as described for the electrophysiology experiments (see above). Slices were fixed in 4% paraformaldehyde in 0.1M phosphate buffer (PB, pH 7.2) at 4°C overnight and then cryoprotected in 20% sucrose in 0.1M PB for 1h prior to subsectioning. For visualization of transfected cholinergic neurons in the septum, deeply anesthetized mice were perfused with 0.9% saline followed by 4% paraformaldehyde in 0.1 M phosphate buffer (PB; pH 7.2) and then cryoprotected overnight in 20% sucrose. In both cases, slices and brains were sub-sectioned (30 μm thickness) on a freezing microtome and sections were mounted onto slides for subsequent processing for immunofluorescence. For visualization of GFP/EYFP expression, fluorescence was enhanced by immunolabeling with chicken anti-GFP (1:1000; Abcam, # ab13970) followed by goat anti-chicken Alexa Fluor 488 (1:1000; Thermo Fisher Scientific, # A-11039).^46^ Tissue sections were coverslipped with Vectashield Mounting Medium containing DAPI (Vector Laboratories, # H-1200-10) prior to imaging. Images were acquired using a Leica DM6000 epifluorescence microscope equipped with a Hamamatsu ORCA-ER digital camera Images and Volocity (V4.0, PerkinElmer) software.

## Results

### Characteristics of MHb neurons

Coronal, ex vivo slices (**Fig 1a**) were used to investigate the effects of cocaine on activity in the ventral portion of the medial habenula (vMHb), a subfield that contains a dense concentration of cholinergic neurons (**Fig 1b**). The cells have thin axons with sizeable terminals that appear to contact other ACh neurons (**Fig 1c**). This is in accord with Golgi studies that describe recurrent collaterals within the vMHb.^47^ The nucleus is innervated by the medial septum / diagonal bands of Broca complex (MS / DBB), a region that contains sizeable populations of GABAergic and cholinergic cells. There has been some uncertainty regarding the MHb receiving input from MS/DBB cholinergic neurons but recent work argues against such a connection.^48^ We found that selective expression of the fluorescent-tagged ChannelRhodopsin 2 (ChR2) construct by choline acetyl transferase (ChAT)-positive cells in the MS/DBB produced dense labeling of fibers and terminals in known targets of the complex (e.g., the hippocampus) but very few labeled processes in the MHb including it’s ventral portion (**Fig 1d**). Multiple lines of evidence indicate that the GABAergic cells in the MS/DBB send axons to the MHb via the stria terminalis.^49^ Other work showed that the GABAergic input to the nucleus is excitatory, an effect that is related to a lack of potassium co-transporter KCC2.^50-52^ We recorded from vMHb neurons using cell attached pipettes, in some cases under visual control for neurons expressing ChAT (**Fig 1e**), and confirmed that stimulation of the stria medullaris triggers spikes. Specifically, a brief train of pulses delivered at 20Hz elicited time-locked discharges from single neurons. The induced spiking was followed by a silent period lasting for a second or more and then a longer interval with a less than normal firing rate **(Fig 1f)**. As expected, single pulse activation of stria medullaris elicited a robust EPSC in vMHb neurons that was blocked by the GABA_A_R antagonist bicuculline (**Fig 1g**). In all, cells recorded in the slice preparations exhibited a diverse array of properties known to characterize the ventral, cholinergic segment of the MHb.

**Figure 1.**
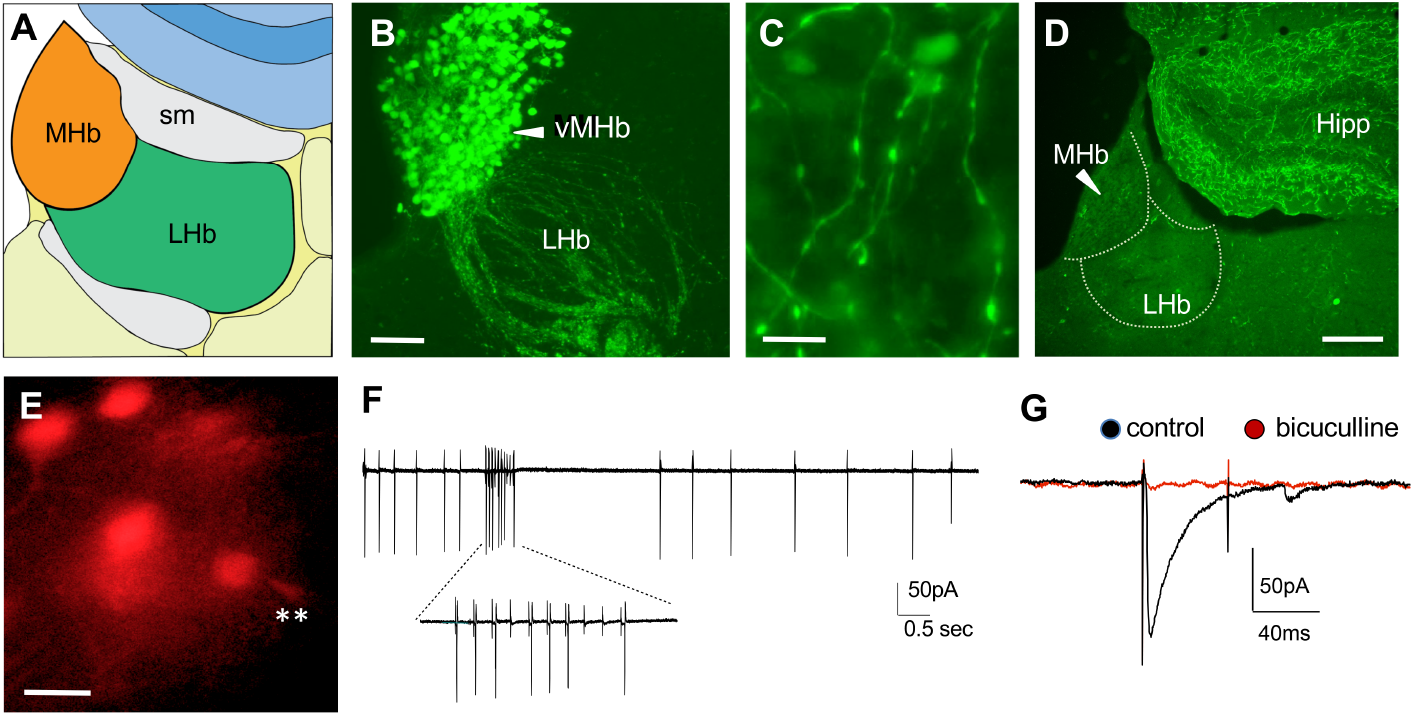
Prominent features of ventral division of the medial habenula (vMHb): **A**) Locations for the two major subdivisions of the habenula, the medial habenula (MHb) and lateral habenula (LHb); sm, stria medullaris. **B**,**C**) Labeling of vMHb cholinergic neurons and axons in GFP-expressing choline acetyltransferase (ChAT)-cre mice. As shown in Panel B, neurons in the vMHb express dense concentrations of ChAT. In addition, thin cholinergic axons with terminals are seen to ramify within the vMHb (C). **D**) Channel rhodopsin labeled cholinergic axons following a viral injection into the medial septum / diagonal bands. Very few fibers are seen in the habenula while a dense projection is evident in the overlying hippocampal (Hipp) dentate gyrus. **E**,**F**) Loose seal recording from choline acetyltransferase expressing vMHb neurons. Location of a recording pipette is indicated by asterisks. Panel F shows a typical recording from a vMHb neuron before, during, and after delivery of a 20Hz train of stimulation pulses to the stria medullaris. Note that the cell spikes at a regular rate prior to stimulation and is activated during the train (expanded trace below). There is an evident depression of spiking for about 1.5 second after the train. **G**) Whole cell recording of the response of a vMHb neuron to single pule stimulation of the stria medullaris prior to and during infusion of the GABA_A_R antagonist bicuculline (20 µM). Calibration bars, 100 µm (B), 20 µm (C), 200 µm (D), and 20 µm (E).

### Cocaine blocks physostigmine-induced increases in spiking by vMHb neurons

The presence of collaterals with numerous terminals in the cholinergic cell field of the vMHb suggests that the system is capable of robust auto-activation. In accord with recent reports, we found that infusion of the acetylcholinesterase (AChE) inhibitor physostigmine, which increases the concentration of evoked and stochastically released acetylcholine, nearly doubled the firing frequency of cells in the vMHb. This effect was eliminated by mecamylamine hydrochloride (10 µM), a broad-spectrum antagonist of nAChRs. Cocaine infusion (10 µM) had no measurable effect on baseline firing (baseline 4.0 ± 0.4Hz; cocaine: 4.0 ± 0.5Hz) but completely blocked the enhanced spiking produced by physostigmine (**Fig 2a**). Specifically, physostigmine alone doubled firing from 3.9 ± 0.7 to 7.8 ± 0.7Hz (p=0.001, paired t-test) but produced no evident change when infused together with cocaine (4.9 ± 0.8 to 4.5 ± 0.8 Hz, p=0.487). The difference between the AChE inhibitor’s effects in the absence vs. presence of cocaine was highly significant (p=0.001).

**Figure 2.**
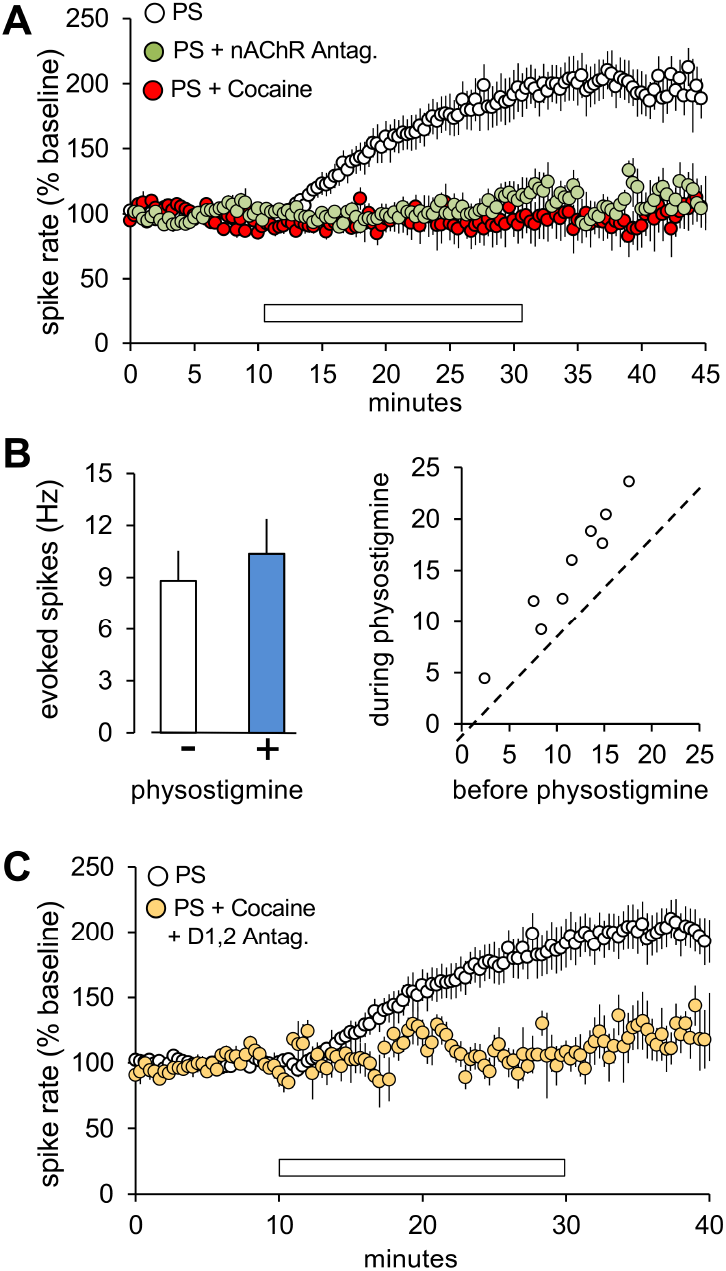
Cocaine blocks ACh driven spiking in the vMHb: **A**) Spikes from single cells were monitored using loose seal recording. An infusion of physostigmine (PS, 10 µM) alone (open bar) caused an approximately two-fold increase in the frequency of action potentials (mean ± s.e.m.; n= 12 cells for PS alone group, n = 7 cells for all other groups). This effect did not occur when the cholinesterase inhibitor was delivered together with either 10 µM cocaine or a broad spectrum antagonist of nicotinic ACh receptors. **B**) (*Left*) Stimulation of the stria medullaris at 20Hz under control conditions elicited 8.8 ± 1.7 within-train spikes above baseline (see Fig 1G). The same stimulation produced 10.4 ± 2.0 spikes above the elevated baseline recorded during physostigmine infusion (n = 9 cells). (*Right*) Slice by slice comparison of evoked spiking recorded before vs. during physostigmine infusion The small deflections from the unity line (dotted line) indicate that stria medullaris input was not synergistic with ACh activation (n = 9 cells). **C**) Infusion of a cocktail of dopamine receptor (D) 1 and 2 antagonists SCH23390 (2.5 µM) and sulpiride (2.5 µM), respectively, did not block the suppression of physostigmine induced spiking by cocaine; drug infusions shown by open bar (n = 5 cells for PS + cocaine + D1,2 Antag. group, 12 cells for PS group).

In a separate set of experiments, we tested for synergy between the increase in spontaneous activity produced by physostigmine and that triggered by stimulation of the stria medullaris. Activation of the input increased the frequency of spiking by 8.8 ± 1.7 Hz during baseline conditions and by 10.4 ± 2.0 Hz when delivered in the presence of physostigmine, an effect that was of borderline significant (p=0.053, paired t-test). In all, evoked activity was greater during than before physostigmine infusion but the magnitude of the effect was small (**Fig 2b, *left***). There was a strong correlation between the increase in spiking produced by 20Hz stimulation of the stria medullaris before vs. during physostigmine infusion (r^2^ = 0.890) but the magnitude of the effect did not exceed 33% (**Fig 2b, *right***). We conclude from this that excitatory GABAergic input does not strongly interact with Ach-driven elevation of baseline firing.

The dramatic inhibitory action of cocaine on Ach-driven activity was surprising because there is little evidence for a strong projection to the vMHb by the dopaminergic systems targeted by the drug.^53, 54^ Moreover, co-treatment with selective antagonists for dopamine D1 (SCH23390 hydrochloride) and D2 (sulpiride) receptors did not interfere with the blocking actions of cocaine (% physostigmine effect from baseline: 4.5 ± 10.4%; p=0.818, paired t-test. (**Fig 2c**). Evidence that cocaine can bind to a subcategory of nicotinic AChRs is of interest with regard to these unexpected findings.

### Induction of cocaine-seeking behavior eliminates subsequent suppression of cholinergic activity by the drug

Past work showed that a brief series of cocaine treatments induces a strong preference for the drug in rodents as shown using conditioned place preference (CPP) and operant learning paradigms. This effect gradually extinguishes over a week of unrewarded trials but is then fully reinstated (‘relapse’) by a single cocaine treatment.^37^ We confirmed these results for the CPP paradigm. Mice were given a habituation period (days 1-3) followed by a pre-test session (day 4) in which they had free access to three rooms connected by open doors. On alternate days (days 5-8) they were injected with cocaine or saline and confined for 30 min in one of two distinctly different end chambers (i.e., the ‘saline’ vs. ‘cocaine’ chamber) (**Fig 3a**). For all test phases, data for each animal was plotted according to their final group assignment on day 15 (i.e., cocaine or saline for reinstatement). The RM-ANOVA showed that within-group differences across test days were highly significant (F(3,42) = 9.5, p<0.0001). There was a strong preference for the cocaine chamber on the day following the alternating cocaine/saline treatments (490.5 ± 16.6 sec for the cocaine chamber vs. 179.9 ± 10.7 sec for the saline chamber) but not after 4 additional days without injections (‘extinction’) (****p<0.0001 for each post-test group vs. their respective behavior pre-test and at extinction, Tukey’s test). An injection of cocaine after extinction (‘cocaine only’ during reinstatement) caused mice to express a strong preference (+193.2 ± 48.8 sec) for the chamber previously associated with the drug (****p<0.0001 vs. their behavior at extinction, and vs. the saline-only reinstatement group, Tukey’s test). Saline treatment did not produce this effect (‘saline only’) (p = 0.5511 vs. their extinction behavior, Tukey’s test). (**Fig 3b**). Next, we tested if direct application of cocaine into the MHb in slices prepared from animals having gone through extinction (no reinstatement) had the same dramatic effects on neuronal firing induced by the local cholinergic system as it did for control slices. The baseline firing rate for vMHb cells in the physostigmine alone group (4.2 ± 0.6Hz) was not detectably different from that described above for untreated mice (3.6 ± 0.4Hz). There was clear tendency for the AChE inhibitor to produce a smaller increase in spiking in the post-extinction group, a point that merits further attention in future studies. In any event, the dramatic effects of infused cocaine on physostigmine-driven increases in neuronal activity recorded from naïve slices were entirely missing in the post-extinction cases (**Fig 3c**). As in the pretreatment slices, cocaine had no effect on baseline (no physostigmine) activity in the extinguished group (pre-infusion spiking rate: 4.1 ± 1.2 Hz; cocaine: 4.4 ± 0.8 Hz). In all, prior experience eliminated the interaction between cocaine and Ach-driven activity in the vMHb.

**Figure 3.**
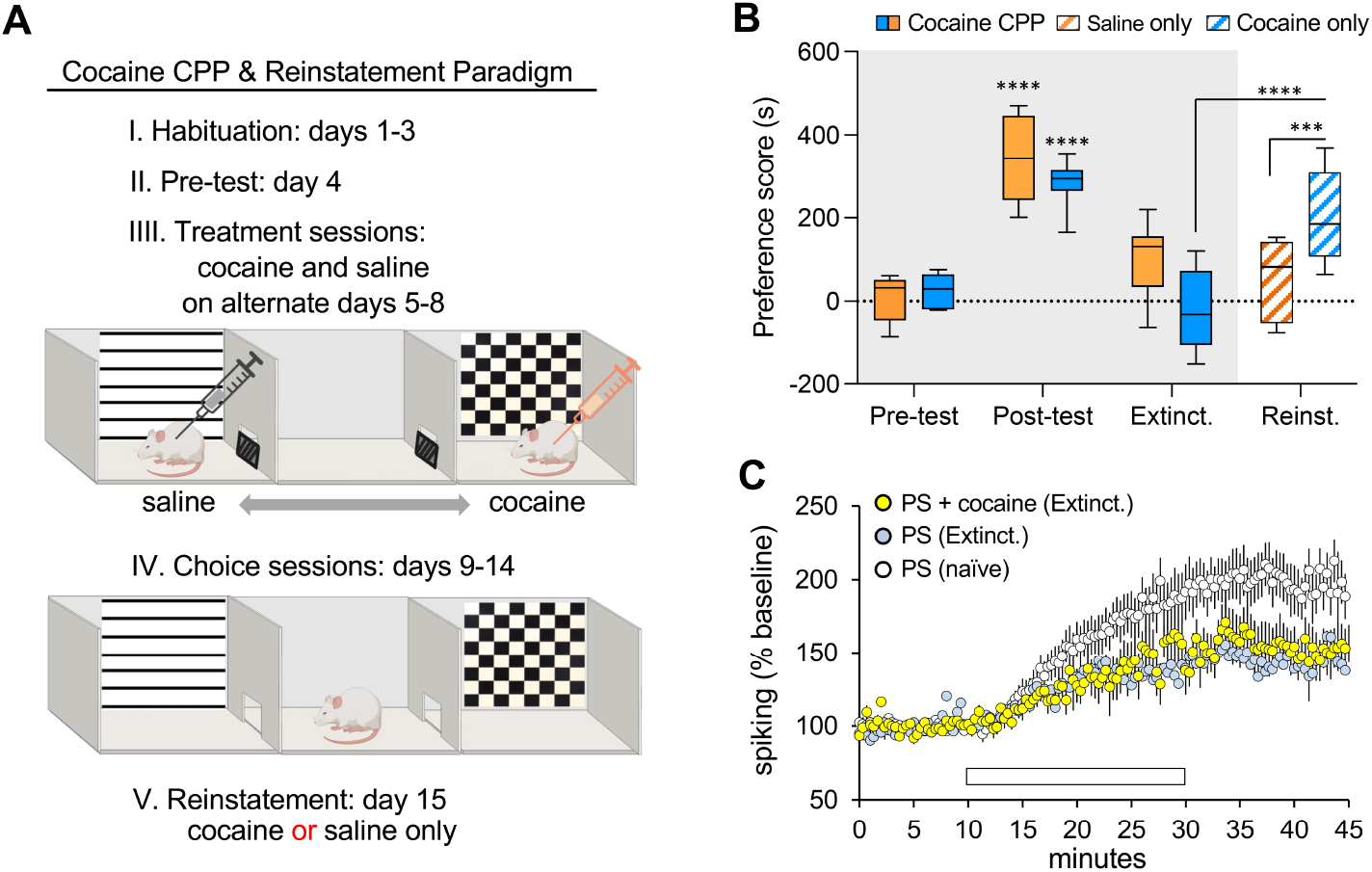
Prior experience with cocaine profoundly changes the effects of cocaine in the vMHb. **A**) Schematic description of the cocaine conditioned place preference (CPP) and reinstatement treatment protocols for behaving mice (see Methods). After habituation (days 1-3) and a pre-test in the 3-chamber apparatus (day 4), all animals received an injection of cocaine (10 mg/kg) before being placed for 30 minutes in one of the end chambers (day 5) with the exit door closed. On day 6, the mice were given a saline injection and placed in the other, visually distinct end chamber. This treatment and chamber pairing were repeated for days 7 (cocaine) and 8 (saline). Starting on day 9 and continuing through day 14, the mice were placed in the central compartment without barriers to the end chambers and allowed to freely explore. For reinstatement on day 15, the mice were randomly assigned to receive either a single injection of cocaine (5 mg/kg) or saline and were again allowed free access to all compartments of the apparatus. **B**) Graph showing the ‘preference score’ (time spent in the cocaine compartment minus time in the saline chamber) for each phase of the cocaine CPP paradigm and following reinstatement on day 15. For all test phases, data for each animal has been plotted according to their final group assignment on day 15 (i.e., cocaine or saline for reinstatement). The preference score for the first free choice day after the chamber specific injections of saline or cocaine was different for the preference scores during ‘pre-test’ or after multiple extinction trials (extinct’) (**** p<0.0001, Tukey test). A single injection of cocaine after extinction restored the preference for the previously established ‘cocaine compartment’ (‘reinst’ vs. ‘extinct’, p<0.0001, Tukey). There was also a statistical difference in preference for the prior ‘cocaine chamber’ in mice given a cocaine vs. saline injection (p<0.001). **C**) Loose seal recordings collected from vMHb slices prepared from mice after the 5^th^ day of extinction; the animals did not receive any further injections of saline or cocaine. The composite results for physostigmine alone treatment in naïve slices are shown for comparison. As shown, cocaine did not reduce the increase in spiking produced by physostigmine in slices prepared from mice that had prior experience with stimulant (n = 7 cells for PS (Extinct.) group, 10 cells for PS + cocaine (Extinct.) group, and 12 cells for PS (naive) group.

## Discussion

The present results describe a novel target for cocaine that has a plausibly direct relationship to the physiological and psychological effects of the drug. The medial habenula is a phylogenetically conserved structure that serves as one of the end stations – the other being the lateral habenula ---for a dorsal diencephalic conduction system that originates in the basal forebrain and ventral pallidum.^55, 56^ Both divisions of the habenula generate dense descending projections via the fasciculus retroflexus to the interpeduncular nucleus and monoaminergic systems of the brainstem.^57-59^ It is widely agreed that these arrangements provide a route whereby limbic and striatal regions regulate arousal, mood, and responses to positive or negative rewards.^60-62^ The nature of this control is still uncertain but considerable evidence suggests that it is predominately inhibitory.^62-65^ If so, then a drug that potently inhibits cholinergic activity in the medial habenula, as shown here for cocaine, would be experienced as rewarding. There are however results suggesting that activation of the cholinergic habenular neurons excites dopaminergic neurons in the VTA.^66^ From this perspective, the observed block of Ach-elicited spiking in the MHb would remove an excitatory input to the VTA and thereby serve as a brake on cocaine’s actions in the striatum.

Cells in the vMHb express high levels of a ‘pacemaker’ channel,^67-69^ resulting in relatively steady rate of baseline firing. Cocaine had no detectable effect on this spontaneous activity at the concentration used here (10 µM). Physostigmine introduced a second type of activity that was entirely blocked by an antagonist of the nAChRs. Given the presence of numerous terminals within the MHb generated by the dense population of resident cholinergic cells, and the absence of extrinsic cholinergic input,^48^ the two-fold increase in spiking produced by physostigmine is indicative of a self-sustained network. Cocaine produced a rapid and complete block of physostigmine-induced cell firing – indeed, it was as effective in this regard as nAChR antagonists. This result provides an example of cocaine producing a dramatic physiological effect that is not mediated by its actions on dopamine reuptake or release.^70^ That is, dopaminergic terminals are few in number in the vMHb^53, 54^ and we found that dopamine receptor antagonists did not counteract the inhibitory effects of cocaine on physostigmine-induced activity. Accordingly, we conclude that in the vMHb cocaine exerts it suppressive effects via previously described binding to nicotinic receptors.

We had expected to record a degree of synergism between ACh-driven cell spiking and the excitatory GABAergic input from stria medullaris but instead found that the latter simply added to the effects of physostigmine. Whether there are stria medullaris stimulation patterns that engage the cholinergic feedback system and extended spiking is an important issue for future research and one that relates to cocaine’s effects on the operations of the vMHb.

In the extinguished mice, prior experience with cocaine may have reduced ACh-induced spiking although the variability in the physostigmine-alone groups in the present studies precludes a firm conclusion. Baseline firing was unaffected by the earlier treatments with the drug, suggesting that any defects produced by the induced preference / extinction protocol likely involved some aspect of cholinergic transmission. The vMHb is known to express dense concentrations of nAChRs^25, 26^ along with high levels of extrasynaptic receptors.^71^ There is also evidence that the cholinergic neurons in the MHb utilize a diverse array of nAChR subunits,^25^ and exhibit experimentally induced changes in expression of those receptors.^72^ Collectively, these observations raise the possibility that cocaine pretreatment altered the expression of nAChRs or the balance of subunit expression and thereby affected the response to released acetylcholine.

Prior experience with cocaine eliminated the acute effects of the drug on physostigmine-induced changes in cell firing. This dramatic and entirely unexpected consequence of inducing (and then extinguishing) drug-seeking behavior could reflect a shift in the presence of nAChR subunit targets of cocaine. Recent studies have identified nAChR subunit combinations that provide binding sites for cocaine ^8-10, 30^ but further work is needed to clarify the relative affinities for receptor variants in the vMHb. It is accordingly not possible to develop a hypothesis with specific predictions about genomic changes. The loss of the cocaine response is also intriguing in light of work showing that a single injection fully reinstates cocaine-seeking behavior (‘relapse’) in extinguished mice. ^37^ The treatment activates the vMHb ^37^ and initiates rapid and pronounced epigenetic changes including a break in the linkage between the histone deacetylase HDAC3 and the transcription factor NR4A2 (a.k.a, NURR1).^42^ The latter is expressed at high levels in the vMHb. Loss of function experiments showed that expression of NR4A2 within the MHb is critical for the long lasting propensity to relapse that follows repeated use of cocaine.^41, 42^ It is somewhat paradoxical that peripherally administered cocaine activates the vMHb and triggers behaviorally critical genomic changes at a time in which the drug has lost its acute effects on physiology within the same region. One possibility is that *in vivo* cocaine acts on the MS / DBB complex to increase the flow of GABAergic input to the vMHb ^49^ to produce dramatic shifts in physiological activity, and that this interacts with an altered state of the local cholinergic cells. It will be of interest in this regard to test if repetitive stimulation of the stria medullaris in slices from previously cocaine treated/extinguished mice elicits the types of genomic effects found after an *in vivo* cocaine treatment.

In all, we propose a three stage model for the consequences of cocaine usage for the operation of the vMHb: 1) first experience: a drug-induced depression of cholinergic activity that contributes to acute psychological responses by altering dopamine activity; 2) renewed use after abstinence (immediate): cocaine does not *acutely* affect recurrent cholinergic activity; 3) renewed use after abstinence (persistent): rapid induction of epigenetic effects that produce a long lasting tendency to relapse to drug–seeking behavior. Whether the adjustments associated with steps 1 and 2 prime the system for the enduring step 3 changes is at this point an open question.

